# BIK1-RIN4 complex links plant immunity and pathogen virulence

**DOI:** 10.1101/2025.09.03.673898

**Authors:** Huiwon Lee, Dohee Ko, Hobin Kang, Sang Hee Kim, Farid El-Kasmi, Jeffery L. Dangl, Eui-Hwan Chung

## Abstract

Plants deploy receptor-like cytoplasmic kinases (RLCKs) and the immune hub protein RIN4 to coordinate PAMPs-triggered immunity (PTI). Here, we show that BIK1, but not its close homolog PBL1, dynamically strengthens the interaction with RIN4 upon flg22 perception, thereby promoting phosphorylation of RIN4 at S141 and potentiating PTI responses. We further demonstrate that the bacterial type III effector AvrB disrupts the BIK1-RIN4 association through phosphorylation of RIN4 at T166, abolishing complex formation and attenuating BIK1-RIN4 complex-dependent PTI. These findings newly elucidate the BIK1-RIN4 complex as a convergent node of plant immunity and effector targeting, providing insight into the molecular arms race between host defenses and pathogen virulence strategies.

---

Plants have developed a two-tiered innate immune system to counter pathogenic microorganisms (Jones et al., 2024). At the cell-surface, pattern recognition receptors (PRRs) perceive pathogen-associated molecular patterns (PAMPs), thereby initiating PAMP-triggered immunity (PTI), which is characterized by early defense responses such as calcium influx, reactive oxygen species (ROS) production, and cell wall fortification (Zipfel & Robatzek, 2010). To suppress PTI, pathogens secrete effector proteins into host cells, leading to effector-triggered susceptibility (ETS). Plants, in turn, utilize nucleotide-binding leucine-rich repeat receptors (NLRs) to recognize these effectors and activate effector-triggered immunity (ETI), characterized by hypersensitive response (Iakovidis et al., 2023).

Receptor-like cytoplasmic kinases (RLCKs) are involved in various plant signaling pathways, with subfamily VII members playing pivotal roles in innate immune responses downstream of PRR recognition during PTI responses (Liang & Zhou, 2018). Genetic and molecular studies have revealed that AvrPphB-susceptible 1 (PBS1)-like proteins (PBLs), including Botrytis-induced kinase 1 (BIK1) and PBS1-like 1 (PBL1), serve as essential integrators of PRR-mediated immune signaling and are critical for robust PTI activation (Zhang et al., 2010). Upon PAMP recognition, PRRs assemble complexes with co-receptor BAK1, which initiate trans-phosphorylation events (Chinchilla et al., 2007). The activated PRR-BAK1 complex subsequently phosphorylates RLCKs such as BIK1 and PBL1, leading to their dissociation from the PRR-BAK1 complex and enabling them to mediate downstream PTI signaling cascades (Zhang et al., 2010).

RPM1-interacting protein 4 (RIN4) is an intrinsically disordered “phospho-switch” hub protein in plant immunity that modulates both PTI and ETI (Chung et al., 2014). RIN4 acts primarily as a negative regulator of PTI. The flagellin-derived peptide flg22 activates flagellin sensing 2 (FLS2)-mediated immune responses and induces phosphorylation of RIN4 at serine 141 (pS141), which positively regulates PTI. However, the host kinase(s) responsible for RIN4 pS141 remain unidentified. Conversely, multiple bacterial effectors (e.g., AvrB, AvrRpm1, AvrRpt2, AvrPphB, HopF2, HopZ5, and AvrBsT) directly target RIN4 and induce post-translational modification to provoke susceptibility (Kim et al., 2023). Notably, *Pseudomonas syringae* effector AvrB mediates phosphorylation of RIN4 at threonine 166 (pT166) through a RLCK RPM1-induced protein kinase (RIPK), and pT166 is specifically required for activation of the NLR immune receptor RPM1 (Chung et al., 2011; Liu et al., 2011).

A previous study using Arabidopsis *bik1pbl1* double mutants revealed reduced phosphorylation of RIN4 S141 upon flg22 treatment (Chung et al., 2014). Phylogenetic analysis indicated that BIK1 and PBL1 are the closest members within the RLCK PBL family (Figure S1A). Despite this close relationship, the two single mutants exhibited contrasting phenotypes: *bik1* showed a dwarf phenotype, whereas *pbl1* was phenotypically indistinguishable from Col-0 (Figure S1B), suggesting their overlapping yet distinct biological functions. Bimolecular fluorescence assays (BiFC) confirmed direct interactions between RLCKs and RIN4 at the plasma membrane (Figure 1A), and subcellular fractionation assays further supported the membrane localization of BIK1, PBL1, and RIN4 (Figure S2). *In vitro* kinase assays using [λ-^32^P] ATP demonstrated that both BIK1 and PBL1 directly phosphorylate RIN4 (Figure 1B). In addition, semi-*in vivo* kinase assays using RIN4 recombinant protein and plant extracts from Col-0, *bik1, pbl1* further supported the contributions of BIK1 and PBL1 to RIN4 phosphorylation (Figure S3). Collectively, these findings provide molecular evidence linking BIK1 and PBL1 to the regulation of RIN4.

**Figure 1.**
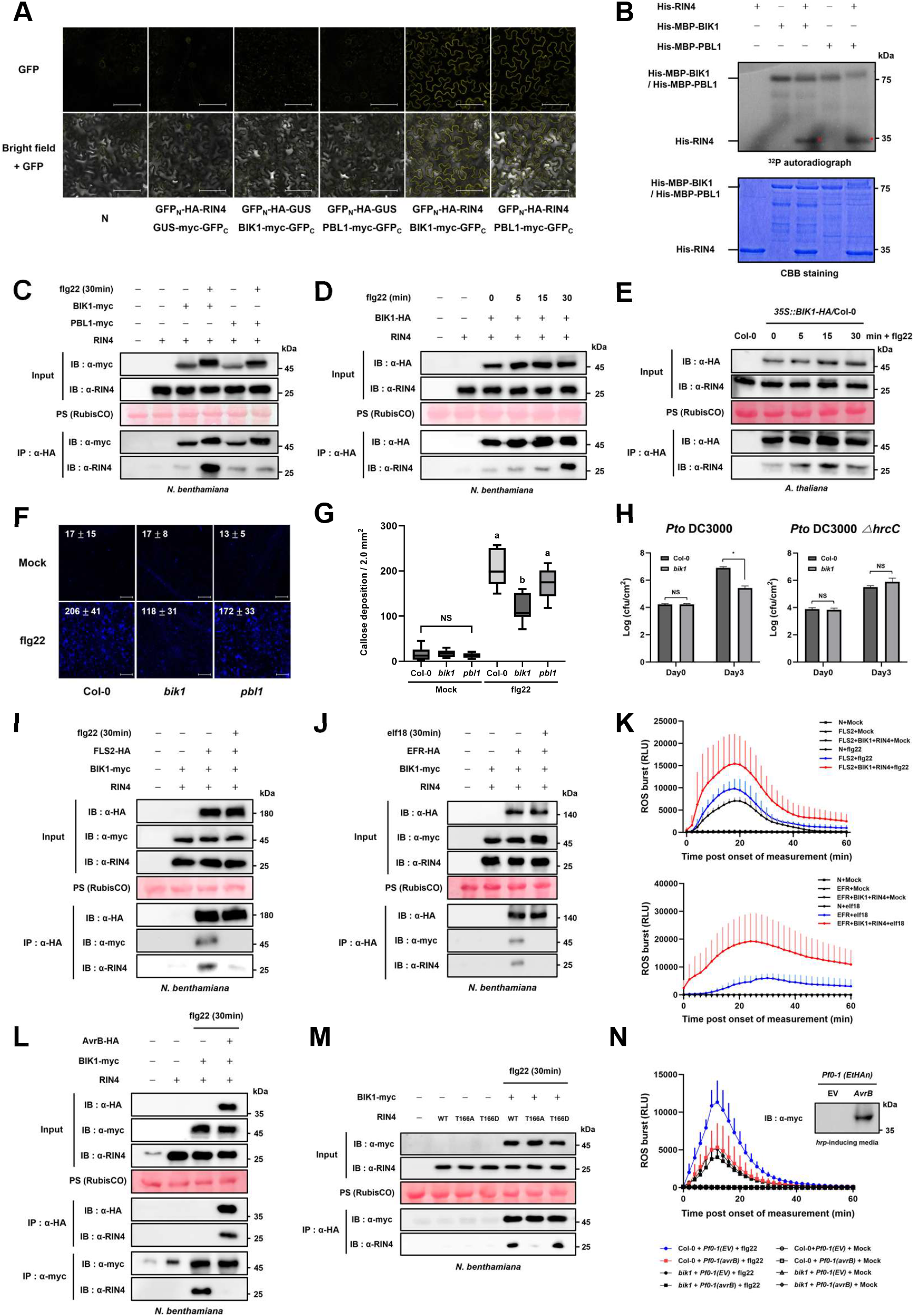
Functional characterization of RLCK-RIN4 complex and bacterial type III effector AvrB during plant immune response. **(A)** Bi-molecular complementation (BiFC) assay to demonstrate the direct interaction between RLCKs and RIN4. GUS construct was used as a negative control. Scale bar, 100 μm. **(B)** *In vitro* kinase assay to confirm the phosphorylation of RIN4 by RLCKs using [λ-^32^P] ATP autoradiography. Red dots indicate phosphorylated RIN4. **(C)** Co-immunoprecipitation (CoIP) analysis to monitor *in planta* interaction between RLCKs and RIN4 with or without flg22-treatment. **(D, E)** CoIP analysis to examine the time-dependent interaction between BIK1 and RIN4 upon flg22-treatment in *N. benthamiana* (D) and *A. thaliana* (E). **(F, G)** Callose accumulation was assessed in Col-0, *bik1*, and *pbl1* after flg22 treatment. Representative images are shown in (F). Values are means ± SE, n=8. Scale bar, 100 μm. Quantification data is presented in (G) (One-way ANOVA followed by Tukey’s multiple range test; different letters indicate significant differences). **(H)** Bacterial growth suppression assay with *Pto* DC3000 wild-type and *ΔhrcC* mutant on Col-0 and *bik1* mutant plants (n=4, \**P* < 0.005, Student’s t-test). **(I, J)** CoIP assay to show the dynamics of PRR-BIK1-RIN4 complexes in *N*.*benthamiana*; FLS2-BIK1-RIN4 complex (I), and EFR-BIK1-RIN4 complex (J). **(K)** Reactive oxygen species (ROS) burst assay to validate the reconstitution of PRR complexes in *N*.*benthamiana*; FLS2 (top), and EFR complex (bottom). **(L)** CoIP assay to monitor BIK1-RIN4 complex with or without bacterial type III effector AvrB in *N*.*benthamiana*. **(M)** CoIP assay to examine *In plant*a association between BIK1 and RIN4 phosphorylation mutants related to AvrB. **(N)** ROS burst assay to confirm AvrB virulence function with *Pf0-1* (EV) or *Pf0-1* (*avrB*) in Col-0 and *bik1* upon flg22 treatment. Immunoblot data shows AvrB protein expression in *hrp*-inducing media.

The physical association and modification of RIN4 by RLCKs may serve as critical molecular events in the PTI signaling pathway. To further examine their dynamics, we investigated the RLCKs-RIN4 associations upon PAMP perception. In the resting state, both BIK1 and PBL1 showed weak interactions with RIN4 *in planta*. Upon flg22 treatment, the BIK1-RIN4 interaction markedly increased within 30 minutes, whereas the PBL1-RIN4 interaction remained unchanged (Figure 1C). Time-course analysis in both *Nicotiana benthamiana* (*N. benthamiana*) and *Arabidopsis thaliana* (*A. thaliana*) confirmed a progressive increase in BIK1-RIN4 interaction after flg22 treatment, peaking at 15 min in *A*.*thaliana*, and at 30 min in *N. benthamaiana* (Figure 1D, 1E). In *35S::PBL1-HA* transgenic plants, the PBL1-RIN4 interaction did not increase after flg22 treatment, consistent with results observed in *N*.*benthamaina* reconstruction system (Figure S4). These results imply that both BIK1 and PBL1 interact with RIN4 prior to PTI activation, but only BIK1 strengthens its association with RIN4 upon flg22 treatment, thereby playing additional roles to modulate RIN4 “phospho-switch”.

Functional assays further highlighted distinct roles of BIK1 and PBL1. In callose deposition and ROS burst assays, *bik1* mutants exhibited reduced PTI responses, while *pbl1* mutants showed no significant difference compared with wild type (Figure 1F, 1G, and S5). These results indicate that, despite their sequence similarity, BIK1 and PBL1 act nonredundantly in plant immunity. PBL1 may contribute to basal phosphorylation of RIN4, whereas BIK1 predominantly mediates PTI downstream signaling by enhancing RIN4 pS141 to control PTI positively. To further test the requirement of BIK1 in PTI, we analyzed the *bik1* mutant and *35S::BIK1-HA* transgenic lines. Consistent with a positive regulatory role, ROS burst and callose accumulation were decreased in *bik1*, but enhanced in *35S::BIK1-HA* (Figures S6A and S6B). In response to the hemi-biotrophic bacterial pathogen *Pseudomonas syringae* pv. *tomato* (*Pto* DC3000), *bik1* mutant displayed susceptible phenotypes. However, when challenged with the *Pto* DC3000 *△hrcC* strain lacking a type III secretion system, resistance was comparable between Col-0 and *bik1* (Figure 1F). Together, these results highlight that BIK1 functions as a central regulator of PTI and bacterial type III effectors may repress PTI by specifically targeting BIK1-mediated signaling pathways as part of their virulence strategy. Notably, *bik1* mutants were more resistant to necrotrophic pathogen *Pectobacterium carotovorum* subsp. *carotovorum (PCC)* compared to Col-0, confirming the multifaceted biological functions of BIK1 in plant immunity (Figure S7).

Upon perception of flg22, BIK1 dissociates from PRRs and interacts with downstream targets such as RbohD and calcium channels to undergo PTI responses successfully through phosphorylation of each target (Zhang et al., 2010). Based on this, we inferred that the BIK1-RIN4 complex formation increased upon PTI activation. Thus, the interaction dynamics of the PRR complex were monitored during immune activation. In the resting state, FLS2, BIK1, and RIN4 were associated within a single complex. Following flg22 treatment, however, the release of the BIK1-RIN4 complex from FLS2 complex was observed at 30 minutes (Figure 1I). Moreover, a similar dynamic of BIK1-RIN4 interaction was also observed in the EFR complex (Figure 1J), indicating that this mechanism is conserved across different PRRs during PTI activation. Agro-infiltration of PRRs increased ROS burst, and co-expression of BIK1 and RIN4 further enhanced PTI outcomes, validating the reliability of the transient expression system for reconstructing PTI system in *N. benthamiana* (Figure 1K). Taken together, we concluded that BIK1 and RIN4 are associated with PRRs in the resting state but dissociate upon PTI initiation, leading to enhanced PTI responses.

Bacterial type III effector AvrB has reported to repress PTI by suppressing RIN4 pS141 with through induction of pT166 via RIPK (Liu et al., 2011). With this, we hypothesized that AvrB might target BIK1-RIN4 complex to promote effector-triggered susceptibility (ETS). We examined the effects of AvrB on the BIK1-RIN4 association 30 minutes after flg22 treatment in the presence or absence of AvrB expression in *N. benthamiana*. RIN4 maintained its association with BIK1 in the absence of AvrB, but this interaction was abolished upon AvrB expression (Figure 1L), suggesting that AvrB competitively disrupted the BIK1-RIN4 complex by binding to RIN4. Moreover, the phospho-mimic RIN4 T166D mutant, which mimics the AvrB-induced phosphorylation state, failed to associate with BIK1. By contrast, wild-type RIN4 and the phospho-dead T166A mutant retained their ability to interact with BIK1 (Figure 1M), underscoring the importance of AvrB-mediated T166 phosphorylation in disrupting this interaction. To confirm whether this molecular interference alters immune responses, ROS burst assays were performed in Arabidopsis Col-0 and *bik1* mutant pre-infiltrated with *Pseudomonas fluorescens Pf0-1* strains carrying either an empty vector or an AvrB-expressing construct. In Col-0, *Pf0-1*(*avrB*) strongly suppressed flg22-triggered ROS production compared to *Pf0-1*(EV), indicating that AvrB attenuates PTI. Notably, the degree of ROS suppression by AvrB in Col-0 was comparable to the basal ROS level in the *bik1* mutant. In the *bik1* background, ROS production did not exhibit difference between *Pf0-1*(EV) and *Pf0-1*(*avrB*) pre-infiltrated leaves, suggesting that AvrB suppresses PTI via a BIK1-dependent pathway as part of its virulence activity (Figure 1N). Collectively, these findings demonstrate that AvrB disrupts BIK1-RIN4 interaction through T166 phosphorylation of RIN4, thereby suppressing BIK1-dependent PTI signaling as a virulence mechanism.

Our study reveals the distinct molecular roles of BIK1 and PBL1 in PTI signaling with RIN4. BIK1 functions as a crucial signal transducer, rapidly enhancing interaction with the hub protein RIN4 upon PAMP recognition, and thereby potentiating robust PTI responses. In contrast, PBL1 appears to contribute to the basal phosphorylation status of RIN4. The bacterial effector AvrB exploits this signaling network to dampen host immunity, primarily by targeting RIN4 and disrupting its association with BIK1. Together, these findings highlight BIK1 and RIN4 as key nodes in the plant immune network and underline the evolutionary arms race between plant defense mechanisms and pathogen counter strategies (Figure S8). However, the precise molecular architecture and regulatory dynamics of the BIK1–RIN4 immune complex remain unresolved, underscoring the need for future studies to unravel its role in fine-tuning plant immunity. In addition, further investigation into the coordinated actions of multiple RLCKs that regulate RIN4 phosphorylation will be essential to fully elucidate the complexity of this immune hub.

## ACKNOWLEDGEMENTS

The authors are very thankful to Professor Eun Kyu Oh (Korea University, Seoul, Republic of Korea) for providing *bik1* mutant line for this study. This work was supported by the National Research Foundation of Korea (NRF) grants (RS202500512558(Global Plant Immunity Research Center), RS202500520276, and RS2023NR077220), Korea University Grant, and BK21 FOUR program (E-H.C). The initiation of this research was funded by Howard Hughes Medical Institute (HHMI: J.L.D).

## CONFLICTS OF INTEREST

The authors declare no conflict of interest.

## AUTHOR CONTRIBUTIONS

E-H.C conceived the study, designed the experiments, wrote the manuscript, and provided funding for the study. H.L. performed the experiments, analyzed the data, and wrote the draft manuscript. D.K. and H.K performed the experiments. S-H.K and F.E.K conceived the story and designed the experiments partly. J.L.D funded to initiate the study.

## Notes

### Competing Interest Statement

The authors have declared no competing interest.

## REFERENCES

Chinchilla, D., Zipfel, C., Robatzek, S., Kemmerling, B., Nurnberger, T., Jones, J. D., Felix, G., & Boller, T. (2007). A flagellin-induced complex of the receptor FLS2 and BAK1 initiates plant defence. Nature, 448(7152), 497–500. 10.1038/nature05999

Chung, E. H., da Cunha, L., Wu, A. J., Gao, Z., Cherkis, K., Afzal, A. J., Mackey, D., & Dangl, J. L. (2011). Specific threonine phosphorylation of a host target by two unrelated type III effectors activates a host innate immune receptor in plants. Cell Host Microbe, 9(2), 125–136. 10.1016/j.chom.2011.01.009

Chung, E. H., El-Kasmi, F., He, Y., Loehr, A., & Dangl, J. L. (2014). A plant phosphoswitch platform repeatedly targeted by type III effector proteins regulates the output of both tiers of plant immune receptors. Cell Host Microbe, 16(4), 484–494. 10.1016/j.chom.2014.09.004

Iakovidis, M., Chung, E. H., Saile, S. C., Sauberzweig, E., & El Kasmi, F. (2023). The emerging frontier of plant immunity’s core hubs. FEBS J, 290(13), 3311–3335. 10.1111/febs.16549

Jones, J. D. G., Staskawicz, B. J., & Dangl, J. L. (2024). The plant immune system: From discovery to deployment. Cell, 187(9), 2095–2116. 10.1016/j.cell.2024.03.045

Kim, H., Ahn, Y. J., Lee, H., Chung, E. H., Segonzac, C., & Sohn, K. H. (2023). Diversified host target families mediate convergently evolved effector recognition across plant species. Curr Opin Plant Biol, 74, 102398. 10.1016/j.pbi.2023.102398

Liang, X., & Zhou, J. M. (2018). Receptor-Like Cytoplasmic Kinases: Central Players in Plant Receptor Kinase-Mediated Signaling. Annu Rev Plant Biol, 69, 267–299. 10.1146/annurev-arplant-042817-040540

Liu, J., Elmore, J. M., Lin, Z. J., & Coaker, G. (2011). A receptor-like cytoplasmic kinase phosphorylates the host target RIN4, leading to the activation of a plant innate immune receptor. Cell Host Microbe, 9(2), 137–146. 10.1016/j.chom.2011.01.010

Zhang, J., Li, W., Xiang, T., Liu, Z., Laluk, K., Ding, X., Zou, Y., Gao, M., Zhang, X., Chen, S., Mengiste, T., Zhang, Y., & Zhou, J. M. (2010). Receptor-like cytoplasmic kinases integrate signaling from multiple plant immune receptors and are targeted by a Pseudomonas syringae effector. Cell Host Microbe, 7(4), 290–301. 10.1016/j.chom.2010.03.007

Zipfel, C., & Robatzek, S. (2010). Pathogen-associated molecular pattern-triggered immunity: veni, vidi…? Plant Physiol, 154(2), 551–554. 10.1104/pp.110.161547

