## supplementary materials, methods, and figures for "BIK1-RIN4 complex links plant immunity and pathogen virulence"

### Supplementary Materials and methods

#### Plant materials and growth conditions

*Arabidopsis thaliana* Col-0 accession, *Nicotiana benthamiana* were used in this study. Col-0, mutants, and transgenic plants were sown in Sunshine Mix #4 (Sun Gro Horticulture, USA) mixed with vermiculite and perlite at a 3:1:1 ratio and stratified in the dark at 4℃ for three days. *bik1* mutant is SALK_005291 and *pbl1* mutant is SAIL_1236_D07. The *35S::BIK1-HA*/Col-0 transgenic lines were provided by Dr. Cyril Zipfel (Kadota et al., 2015). The *35S::PBL1-HA*/Col-0 transgenic lines were generated by transforming Col-0 with *35S::PBL1-HA*. Arabidopsis plants were grown at 23 ℃ under short-day conditions (9 h light and 14 h dark). Four or five-week-old plants were used for the experiments. *N. benthamiana* were sown and grown under the same soil conditions as Arabidopsis, at 24 ℃ under 12 h light / 12 h dark conditions. Five-week-old plants were used for the experiments.

#### Plasmid constructions

cDNAs of BIK1, PBL1, and RIN4 were cloned into the pGEN_BsaI vector to generate entry clones using the Golden Gate assembly method. Entry clones were transferred into destination vectors containing epitope tags via Gateway LR reaction (Invitrogen), following the manufacturer's instructions. For *in-vitro* assays, entry clones were introduced into pDest-His-MBP and pTH34 vectors and expressed in *E.coli* strain BL21. For *in-planta* protein expression, entry clones were transferred into pGWB14(*35S::ORF-HA*) and pGWB17 (*35S::ORF-myc*) vectors. For the BiFC assay, cDNAs of GUS, RIN4, and RLCKs were first cloned into pDONR221 donor vector using Gateway BP reaction, and the recombined into a 2-in-1 BiFC destination vector via the LR reaction. RIN4 was fused to the N-terminal YFP fragment at its N-terminus, and RLCKs were fused with C-terminal YFP fragment at the C-terminus (Mehlhorn et al., 2018)

#### ROS burst

Leaf discs from Col-0, *bik1*, *pbl1*, *35S::BIK1-HA*/Col-0, and *35S::PBL1-HA*/Col-0 were placed in white 96-well plates with 200 uL of distilled water in each well. After overnight incubation under light, the water was discarded, and 100 µL of reaction mix-containing 100 nM of flg22, 30 µg/mL of luminol, and 20 µg/mL of horseradish peroxidase (HRP)- was added to each well. Luminescence was measured at 2 min intervals for 60 min using Centro luminometer (Berthold technologies, Germany)

**Bacterial growth suppression assay**

Bacterial growth of *Pto* DC3000 wild-type, *ΔhrcC* and *PCC* was assessed to confirm disease resistance in Arabidopsis Col-0 and *bik1*. Bacterial suspensions were prepared in 10mM $MgCl_2$ at $10^6$ CFU/mL and syringe infiltrated. Bacterial growth was quantified 0 and 3 days post-inoculation..

**Agrobacterium-mediated transient assay in *N. benthamiana***

Binary constructs of interest were transformed into *Agrobacteria tumefaciens* strain GV3101. The bacteria were grown overnight in liquid LB medium at 28°C, then pelleted by centrifugation at 13,000 rpm for 2 min at 4°C. The pellets were resuspended in induction media (10 mM $MgCl_2$, 10mM MES pH 5.6, 0.2 mM acetosyringone (Sigma-Aldrich)) and incubated at room temperature for 2 h. The bacterial suspensions were then adjusted to appropriate concentration for each construct and infiltrated into *N. benthamiana* leaves using a 1 mL needleless syringe. Leaf samples were collected 2 days post-infiltration for immunoblotting analysis.

**BiFC (Bi-molecular Fluorescence Complementation) assay**

BiFC assay was performed with Agrobacteria-mediated transient expression *in N. benthamiana*. Plasmid constructs were prepared as described above. Agrobacterium strains harboring the 2-in-1 BiFC constructs were adjusted to an $OD_{600}$ of 0.3 and syringe infiltrated. Two days post-infiltration, leaf tissues were imaged using an LSM-700 confocal microscope (Zeiss). Fluorescence signals were detected using GFP filter settings with an excitation wavelength of 485 nm.

**Protein extraction and immunoblot**

Plant tissue samples from *A. thaliana* and *N. benthamiana* inoculated with binary vectors of interest were harvested in liquid nitrogen and ground in protein extraction buffer (20mM Tris-HCl pH 7.5, 150mM NaCl, 1% Triton X-100, 1mM EDTA pH 8.0, and 0.1% SDS), supplemented with 10mM DTT and 1x plant protease inhibitor cocktail (Sigma-Aldrich). Extracts were centrifuged at 14,000 rpm for 10 min at 4°C, and supernatants were carefully collected, and protein concentrations were determined using the Bradford assay (Bio-Rad). Equal amounts of protein were mixed with 6X SDS sample buffer (0.5 M Tris pH 6.8, 50% glycerol, 10% SDS, 0.1% bromophenol blue, 0.05% β-mercaptoethanol) and boiled at 95 °C for 3 min prior to SDS-PAGE. Protein samples were separated by SDS-PAGE and transferred to a PVDF membrane (GE Healthcare). Immunoblotting was performed using α-HA (Roche, 1:3000), α-myc 9B11 (Cell signaling, 1:2000), and α-RIN4 (Genscript, 1:2000). Detection was carried out using ECL (BIONICS) and visualized with a ChemiDoc imaging system (Bio-Rad).

**Co-immunoprecipitation**

The association of RIN4 with FLS2, BIK1, PBL1, and type III effector AvrB was examined by co-immunoprecipitation. Leaf tissues expressing the proteins of interest were frozen in liquid nitrogen and ground in GTEN buffer (10 % glycerol, 50 mM Tris-HCl pH 7.5, 2 mM EDTA, 150 mM NaCl) supplemented with 5 mM DTT, 1x plant protease inhibitor cocktail (Sigma-Aldrich), and 0.2% NP-40. Protein extracts were centrifuged at 3000 g for 10 min at 4°C. Solubilized supernatants were filtered through Miracloth (G.E. Healthcare). Cleared extracts were incubated with anti-HA- or anti-myc-conjugated magnetic beads (Miltenyi Biotec) for 2 h at 4°C. The beads were washed three times with GTEN buffer, and bound proteins were eluted in 2X SDS sample buffer. Immunoblotting was performed as described above.

***hrp*-inducing protein expression**

*Pf0-1* strains harboring either the AvrB-expressing construct or an empty vector were cultured overnight in liquid KB medium at 28°C. Cells were collected by centrifugation at 9,000 rpm for 6 min at 4°C and resuspended in *hrp*-inducing minimal medium (50 mM potassium phosphate, 7.6mM $(NH_4)_2SO_4$, 1.7mM $MgCl_2$, 1.7mM NaCl, 10mM fructose) at $OD_{600}$ 0.4. After incubation for 6 hours, cells and supernatant were separated by centrifugation at 13,000 rpm for 2 min at 4°C. Cell pellets were resuspended with 2X SDS sample buffer, and immunoblotting was performed as described above.

***In-vitro* kinase assay**

~2 µg of each protein was mixed in kinase buffer (200 mM Hepes pH 7.5, 200 mM MgCl2, 20mM MnCl2, and 50 µM [γ-32P] ATP) of 20 µL. The 4 µCi [γ-32P] ATP was applied in the reaction. Subsequently, the reaction was performed at 30℃ for 30 min and stopped by adding 5x SDS-loading buffer. The phosphorylation status was visualized by autoradiography after loading it on the SDS-PAGE gel.

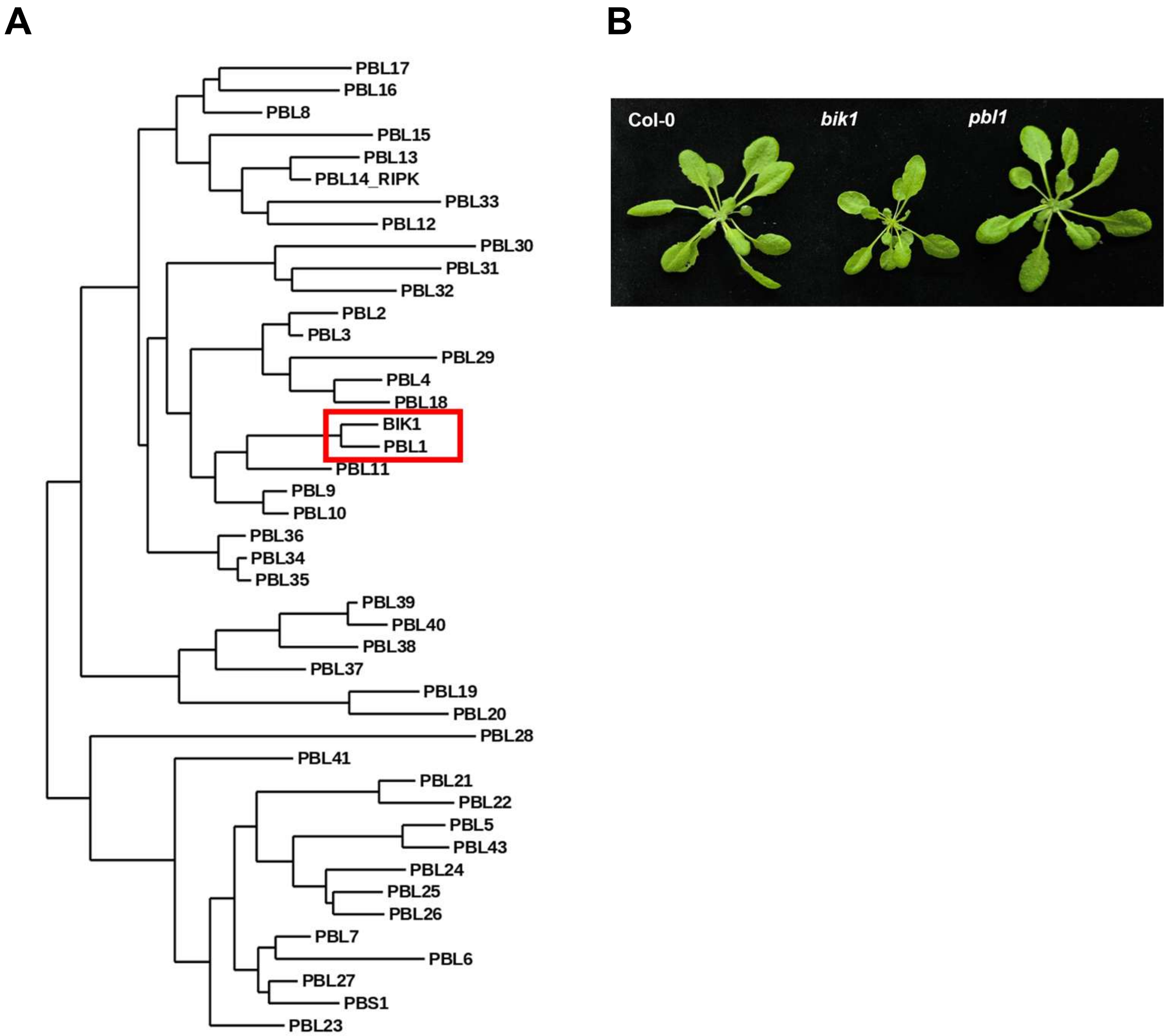


**Figure S1. BIK1 and PBL1 are closely related but distinct kinases.**

**(A)** Phylogenetic tree analysis of receptor-like cytoplasmic kinase (RLCK) PBL (PBS1-like) family members. BIK1 and PBL1 are closely clustered together, which is indicated by a red box. **(B)** Phenotype of *A. thaliana* Col-0, *bik1*, and *pbl1*. Five-week old plants were pictured.

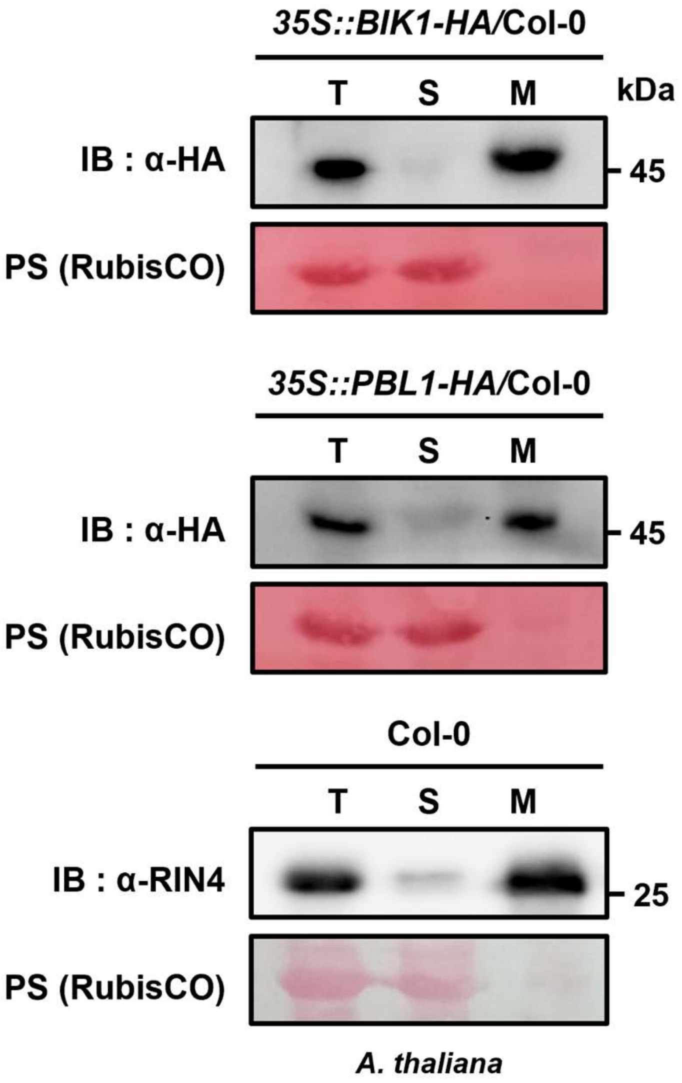


**Figure S2. Plasma membrane localization of BIK1, PBL1, and RIN4.**

Subcellular fractionation assay with *A.thalina* Col-0 and transgenic plants. Protein extracts were separated into total (T), soluble (S), and microsomal (M) fractions. Ponceau S staining of RubisCO served as a loading control, verifying the reliability of the fractionation procedure.

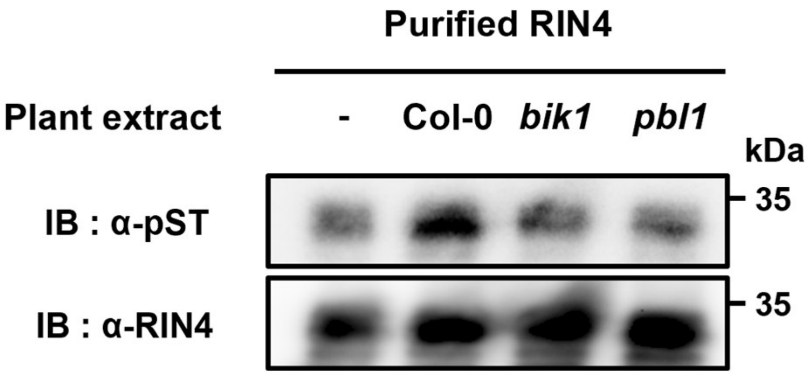


**Figure S3. BIK1 and PBL1 are sufficient to induce phosphorylation of RIN4.**

2 μg of recombinant His-RIN4 protein was incubated with 5 μg of total plant extracts from *A. thaliana* Col-0, *bik1,* or *pbl1* in a semi-*in vivo* kinase assay. Phosphorylated RIN4 was detected by immunoblotting with a anti-phospho-serine/threonine antibody (α-pST), and equal loading was confirmed with an anti-RIN4 antibody (α-RIN4).

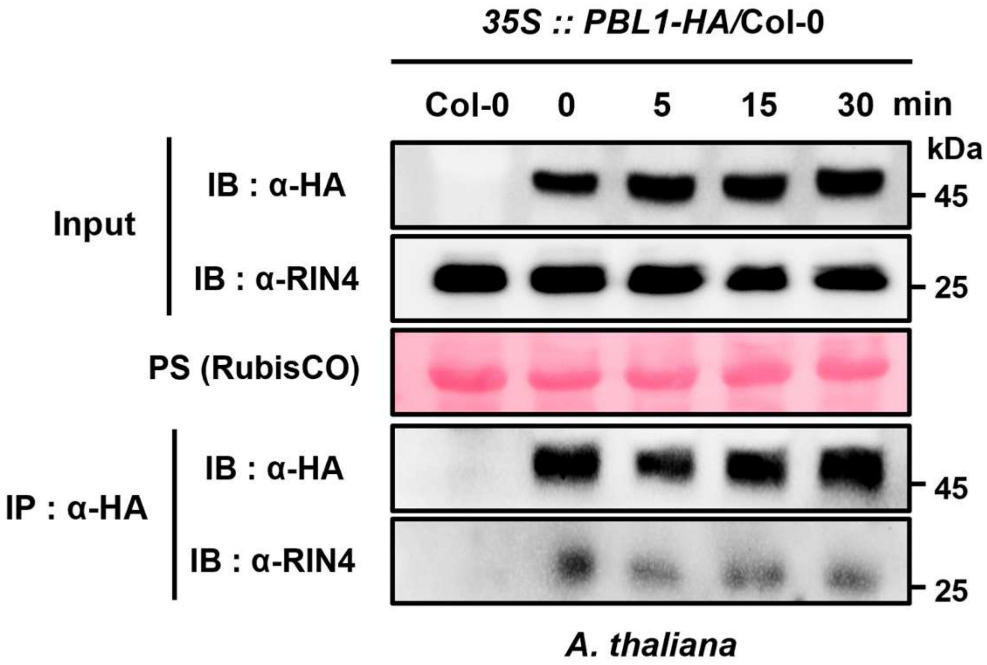


**Figure S4. Constant interaction between PBL1 and RIN4 upon flg22 treatment.**

*35S::PBL1-HA/*Col-0 transgenic lines were infiltrated with 100 nM flg22. Samples were collected at each indicated time point after flg22 treatment. Co-immunoprecipitation was performed with α-HA magnetic beads to confirm the physical association. Immunoblots with α-HA and α-RIN4 confirmed the expression of BIK1 and RIN4.

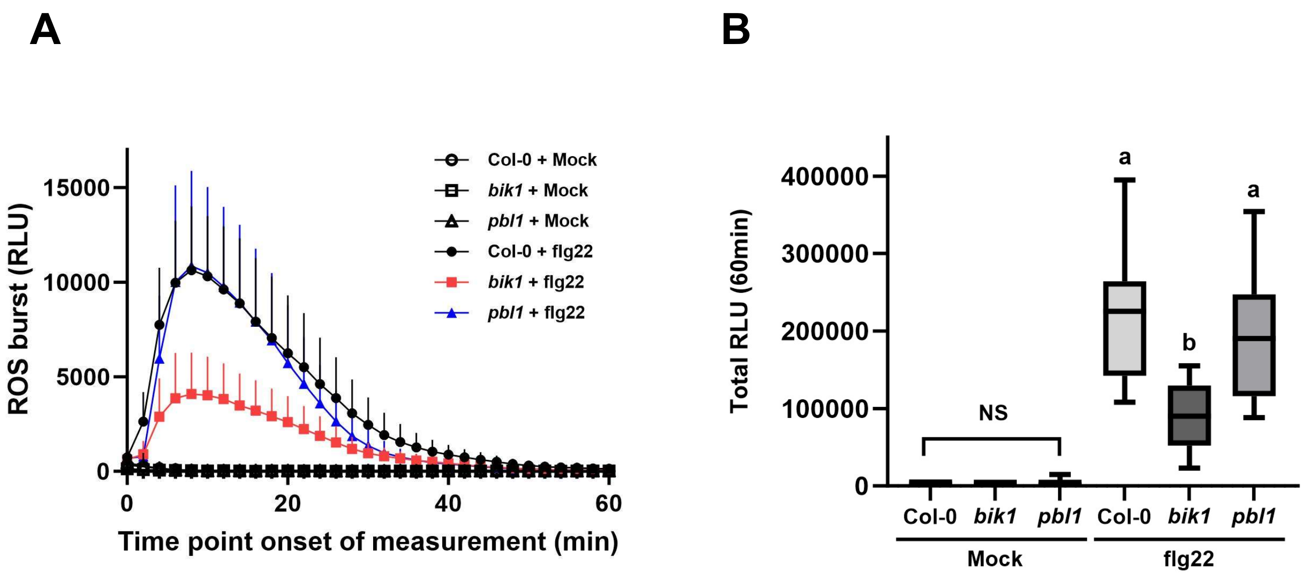


**Figure S5*. bik1* showed reduced ROS production.**

**(A)** Reactive oxygen species (ROS) burst of *A. thaliana* Col-0, *bik1*, and *pbl1*. 100nM flg22 was treated and the ROS production was measured over 60 min. **(B)** Total ROS production for 60 min was quantified, and statistical significance among groups was determined by one-way analysis of variance (ANOVA) followed by Tukey's multiple comparison test; different letters indicate significant differences.

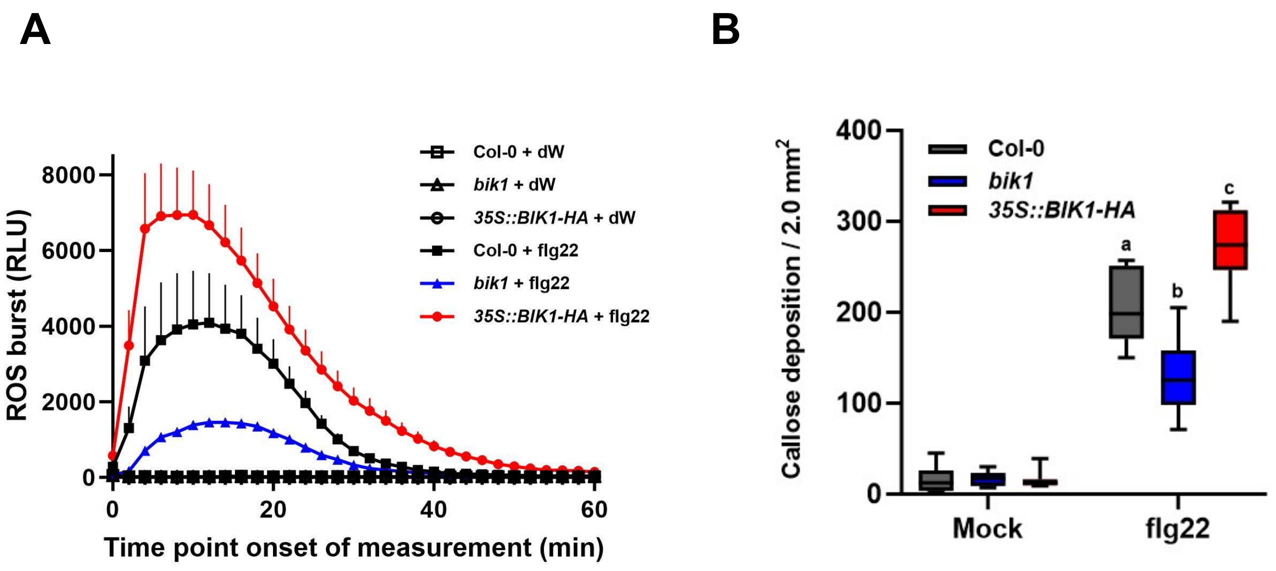


**Figure S6*. BIK1 is a critical integrator of PTI signaling pathway*.**

**(A)** Reactive oxygen species (ROS) burst of *A. thaliana* Col-0, *bik1*, and *35S::BIK1-HA*/Col-0. 100nM flg22 was treated and the ROS production was measured over 60 min. **(B)** Callose accumulation was assessed in Col-0, *bik1*, and *35S::BIK1-HA*/Col-0 after flg22 treatment (n=8). Statistical differences among groups were evaluated using one-way ANOVA followed by Tukey's multiple comparison test; different letters indicate significant differences.

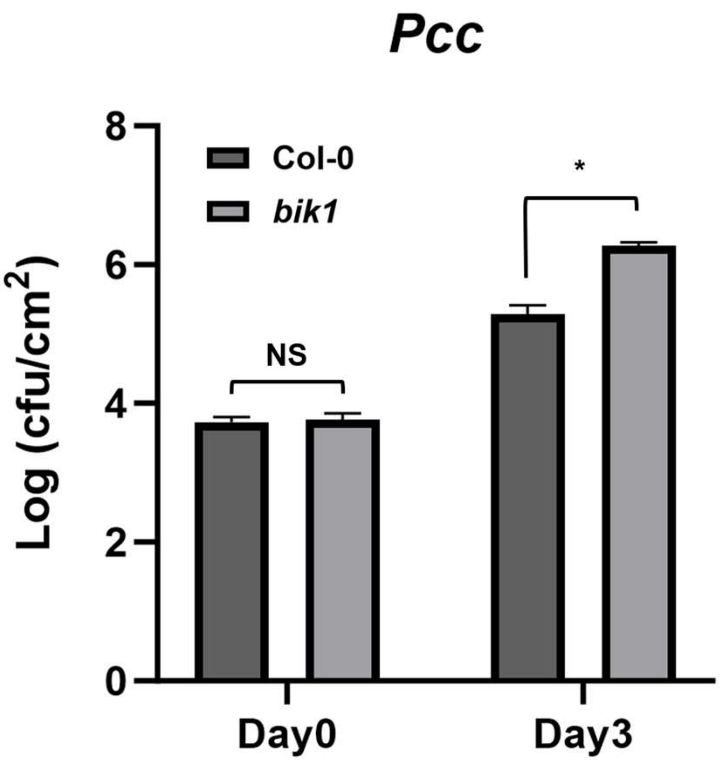


**Figure S7. BIK1 compromises disease resistance against a necrotrophic bacterial pathogen**

Bacterial growth suppression assay of *Pectobacterium carotovorum* subsp*. carotovorum (PCC)* on Col-0 and *bik1*. Bacterial suspension at $OD_{600}$=0.002 was syringe-infiltrated into leaves (n=4). Bacterial growth was quantified, and statistical significance was assessed using Student's t-test (**P* < 0.005).

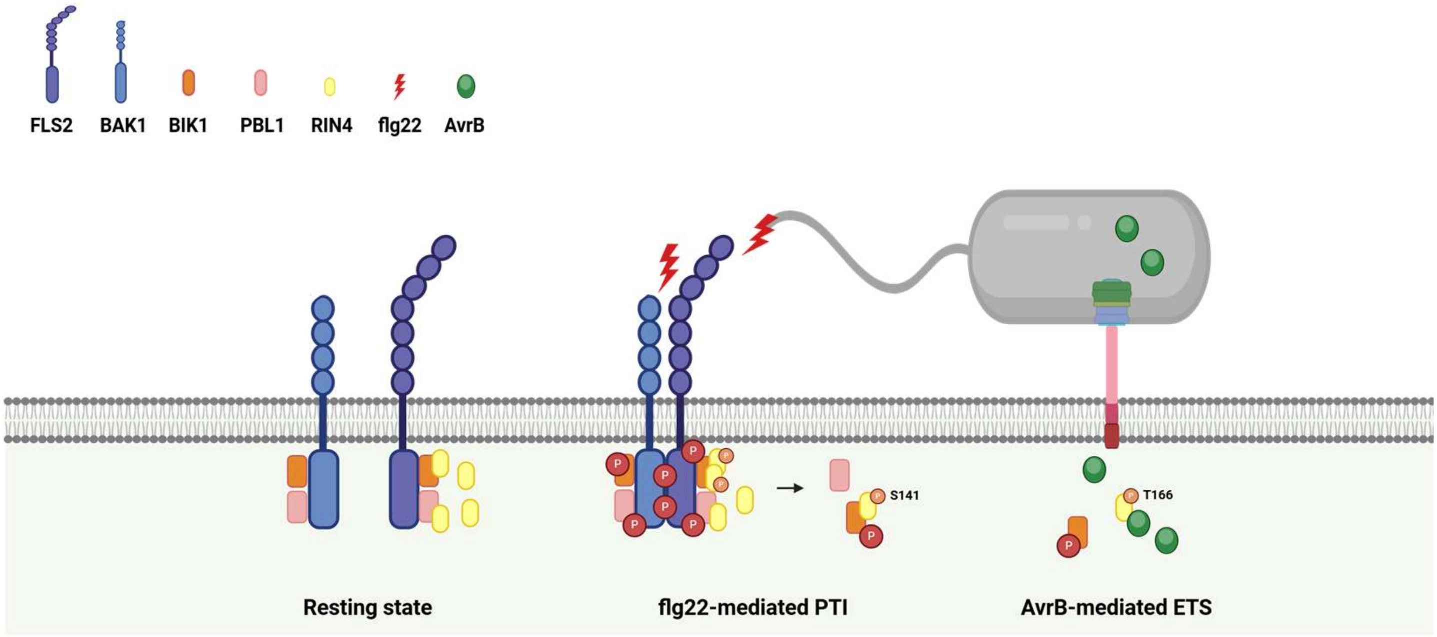


**Figure S8. Proposed working model**

In the resting state, PRRs, RLCKs, and RIN4 form a single protein complex at the plasma membrane. Upon PAMPs recognition, PRR recruits co-receptor BAK1, and the BIK-RIN4 complex dissociates from PRRs, enhancing their association to mediate downstream PTI signaling, including phosphorylation of RIN4 at S141. The bacterial type III effector AvrB disrupts the BIK1-RIN4 complex to promote susceptibility. Together, BIK1-RIN4 is a critical functional module in PTI, specifically targeted by AvrB as its virulence mechanism.

| Primer | Sequence (5'-3') | Description |
|---|---|---|
| BIK1_entry-F | AGTCGGTCTCACTTCATGGGTTCTTGCTTCAGTTCTCGA | Entry clone |
| BIK1_entry-R (Stop O) | AGTCggtctcAGGTGCTACAATCCAACGGTTTTTTTGTTTAAACCG | Entry clone |
| BIK1_entry-R (Stop X) | AGTCGGTCTCAGGTGCAATCCAACGGTTTTTTTGTTTAAACCG | Entry clone |
| PBL1_entry-F | AGTCggtctcACTTCATGGGTTCTTGTCTCAGTTCTCGTG | Entry clone |
| PBL1_entry-R (Stop O) | AGTCggtctcAGGTGCTACAATCCAACGGTTTTTTTGTTTAAACCG | Entry clone |
| PBL1_entry-R (Stop X) | AGTCggtctcAGGTGCGCCAATCCAACGGTTTTTTTGTTTAAACCG | Entry clone |
| RIN4_entry-F | AGTCggtctcACTTCATGGCACGTTCGAATGTAC | Entry clone |
| RIN4_entry-R | AGTCggtctcAGGTGTCATTTTCCTCCAAAGCCAAA | Entry clone |
| AvrB_entry-F (GW) | CAAAAAAGCAGGCTCCATGGGCTGCGTCTCGTCAAAAAGCACC ACAG | Entry clone |
| AvrB_entry-R (GW) | AGAAAGCTGGGTGAAAGCAATCAGAATCTAGCAAGCTTCTGTAT TTTTC | Entry clone |
| BiFC_BIK1_entry-F (attB1) | ACAAGTTTGTACAAAAAAGCAGGCTATGGGTTCTTGCTTCAGTT CTCGAG | BiFC entry clone |
| BiFC_BIK1_entry-R (attB4) | CAACTTTGTATAGAAAAGTTGCCGCCACAAGGTGCCTGC | BiFC entry clone |
| BiFC_PBL1_entry-F (attB1) | ACAAGTTTGTACAAAAAAGCAGGCTATGGGTTCTTGTCTCAGTT CTCGTG | BiFC entry clone |
| BiFC_PBL1_entry-R (attB4) | CAACTTTGTATAGAAAAGTTGCCGCCAATCCAACGGTTTTTTTGT TTAAACCG | BiFC entry clone |
| BiFC_RIN4_entry-F (attB3) | CAACTTTGTATAATAAAGTTGCAATGGCACGTTCGAATGTACCAA | BiFC entry clone |
| BiFC_RIN4_entry-R (attB2) | ACCACTTTGTACAAGAAAGCTGGGTTCATTTTCCTCCAAAGCCA AAGCA | BiFC entry clone |
| BiFC_GUS_entry-F (attB1) | ACAAGTTTGTACAAAAAAGCAGGCTatgttacgtcctgtagaaacccca | BiFC entry clone |
| BiFC_GUS_entry-R (attB4) | CAACTTTGTATAGAAAAGTTGCttgtttgcctccctgctgcg | BiFC entry clone |
| BiFC_GUS_entry-F (attB3) | CAACTTTGTATAATAAAGTTGCAatgttacgtcctgtagaaacccca | BiFC entry clone |
| BiFC_GUS_entry-R (attB2) | ACCACTTTGTACAAGAAAGCTGGGTtcattgtttgcctccctgctgc | BiFC entry clone |
| bik1 genotyping -F | CCCCAAATAGTTAATCTCTGTCTGAAACC | Plant genotyping |
| bik1 genotyping –R | CTACAATCCAACGGTTTTTTTGTTTAAACCG | Plant genotyping |
| pbl1 genotyping –F | ATGGGTTCTTGTCTCAGTTCTCGT | Plant genotyping |
| pbl1 genotyping –R | CCCCAAATAGTTAATCTCTGTCTGAAACC | Plant genotyping |

**Table S1. Primers used in this study**
